# Supramammillary projections to the lateral preoptic area drive dopamine release and active behavior

**DOI:** 10.64898/2026.09.01.748391

**Authors:** Yosuke Arima, Zengyou Ye, Coleman B. Calva, Xia Min, John Gibbons, Jorge Mendoza, Beyonce Getachew, Sarah T. Johnson, Satoshi Ikemoto

**Affiliations:** Neurocircuitry of Motivation Section, Behavioral Neuroscience Research Branch, Intramural Research Program, National Institute on Drug Abuse, National Institutes of Health, Baltimore, MD, United States; Division of Movement Disorders, Department of Neurology, Beth Israel Deaconess Medical Center and Harvard Medical School, Boston, MA, United States

**Keywords:** supramammillary nucleus, lateral preoptic area, mesolimbic dopamine, approach behavior, ventral tegmental area, nucleus accumbens

## Abstract

The supramammillary nucleus (SuM) is known to promote active, approach-oriented behaviors by engaging ventral tegmental area (VTA) dopamine neurons projecting to the nucleus accumbens (NAc), primarily via a glutamatergic SuM→medial septum (MS) pathway. However, the role of other SuM projections in this process remains unclear. Here we combined anatomical tracing, fiber photometry, chemogenetic, and optogenetic approaches to investigate the SuM→lateral preoptic area (LPO) pathway. We found that SuM→LPO neurons possess extensive collateral projections and are inhibited during consummatory behavior but activated by aversive and unexpected stimuli. Chemogenetic activation of these neurons reduced water intake and immobility during stress, while inhibition enhanced the appetitive phase for ingestive behavior and fear-related immobility. Optogenetic stimulation of SuM→LPO terminals reinforced behavior and elicited dopamine release in the NAc. These findings identify the SuM→LPO pathway as a distinct circuit that, alongside the SuM→MS pathway, engages mesolimbic dopamine signaling to facilitate active behavioral responses to environmental challenges.

**SIGNIFICANCE STATEMENT:** Active coping with threat or opportunity may depend on the supramammillary nucleus (SuM) to mobilize dopamine-driven approach behavior. This study identifies the lateral preoptic area as a second SuM target capable of driving dopamine release and reinforcing approach behavior, with activity that tracks whether an animal is facing a threat or consuming a reward. Because SuM neurons broadcast to this and other targets through extensive collateral branches, these results reshape how the SuM’s role in active, approach-oriented behavior should be understood — not as a single circuit, but as a distributed one.

## INTRODUCTION

### The Supramammillary Region and Approach Behaviors

The supramammillary nucleus (SuM) plays a critical role in promoting active, approach-oriented behaviors in response to environmental challenges (Kesner, 2022). SuM neurons respond robustly to salient events, including stress, novelty, and anticipation of food (Beck and Fibiger, 1995; Wirtshafter et al., 1998; Le May et al., 2019; Zhang et al., 2026). Rather than mediating passive defensive behaviors such as freezing or grooming, activation of SuM neurons elicits vigorous, active responses such as jumping, digging, treading, and struggling under restraint (Escobedo, 2024).

These active responses may depend on the SuM’s ability to engage the mesolimbic dopamine system (Kesner et al., 2021). Stimulation of SuM neurons activates dopamine projections from the ventral tegmental area (VTA) to the nucleus accumbens (NAc) and reinforces instrumental approach behaviors (Ikemoto; Ikemoto et al., 2004; Ikemoto, 2005; Ikemoto et al., 2006; Kesner et al., 2021). Collectively, these findings suggest that SuM networks mobilize active actions during threat or uncertainty by recruiting downstream mesolimbic dopamine pathways (Kesner et al., 2022).

One established pathway through which SuM neurons generate active actions involves glutamatergic SuM neurons projecting to the medial septum (MS), which then excite VTA dopamine neurons (Kesner et al., 2021). Activation of SuM VGluT2 neurons projecting to the MS reinforces approach behavior and activates VTA→NAc dopamine neurons. Thus, SuM→MS glutamatergic neurons are currently considered a key route for regulating active coping responses. However, SuM neurons possess extensive collateral projections (Vertes and McKenna, 2000; Farrell et al., 2021; Escobedo et al., 2024), raising the question of whether additional downstream targets also contribute to the coordination of active responses.

### The Lateral Preoptic Area: A Key Target of SuM Projections

A major downstream target of SuM projections is the lateral preoptic area (LPO) (Vertes, 1992; Hahn et al., 2022). The LPO is strongly interconnected with the VTA and serves as a critical node for reward-seeking behaviors (Arvanitogiannis et al., 1996; Barker et al., 2017; Reichard et al., 2019; Gordon-Fennell et al., 2020a; Gordon-Fennell et al., 2020b). LPO neurons are activated by aversive stimuli, and direct stimulation of the LPO reinforces instrumental approach behavior and excites VTA dopamine neurons (Gordon-Fennell et al., 2020a). These properties suggest that the LPO is a key component of neural circuits generating active responses to threat or uncertainty, with the SuM as a plausible upstream modulator.

Recent work shows that SuM neurons projecting to the broader preoptic area are activated by acute stressors, and stimulation of this pathway promotes active responding during stressful conditions; however, activation of this circuit can also induce aversive or avoidance behavior (Escobedo, 2024). These findings raise the important question of whether SuM-driven active responses depend solely on VTA→NAc dopamine neurons, or whether alternative SuM pathways can also generate active behaviors.

### The Present Experiments

To clarify how SuM neurons communicate with downstream regions, the present study addressed four key questions. First, we mapped the extent to which SuM→MS and SuM→LPO neurons possess collateral projections, finding that both populations show extensive collateralization with substantial overlap. Second, we examined SuM→LPO activity during salient positive and negative stimuli, observing that these neurons are activated by threatening events and inhibited during ingestive behavior. Third, we investigated how manipulation of SuM→LPO+Collateral neurons affects ingestive, fear-related, and stress-coping behaviors, finding that activation enhances active coping responses while inhibition promotes passive responses and facilitates the appetitive phase of ingestive behavior. Finally, we tested whether selective SuM→LPO activation is reinforcing and triggers dopamine release, and found that such activation reinforced approach behavior and drove dopamine release in the NAc.

Taken together, these results demonstrate that SuM efferents have extensive collateral organization, enabling them to influence multiple downstream targets in parallel. Both activity recordings and manipulations of SuM→LPO neurons support the conclusion that SuM neurons regulate active responses to environmental challenges. Furthermore, the study identifies the SuM→LPO pathway as an anatomically distinct circuit—in addition to the SuM→MS→VTA dopamine pathway—through which SuM neurons can generate active behaviors.

## METHODS

### Animals

C57BL/6J mice were purchased from the Jackson Laboratory. VGluT2-Cre mice (*Slc17a6*^tm2(cre)Lowl^/J) on a C57BL/6J background were bred at the National Institute on Drug Abuse (NIDA) Transgenic Breeding Facility. Both male and female mice were used in all experiments. Mice were 2–4 months old and weighed 25–35 g at the time of surgery. When not undergoing behavioral testing, mice were group-housed in a temperature– and humidity-controlled vivarium (70–74 °F; 35–55% humidity) under a reversed 12:12 h light-dark cycle with lights off at 07:00. Food and water were available ad libitum, except for mice assigned to water-restriction experiments. For experiments using water as a reward, mice were placed under moderate water restriction for at least 48 h before behavioral testing to enhance water-seeking behavior. During restriction, mice were given access to water for 10 min per day, typically after behavioral testing. Body weight was monitored daily throughout the restriction period.

### Viral vectors, tracers, and pharmacological agents

All AAVs were purchased from Addgene, except DA3m, which was sourced from Biohipp. AAV titers were approximately 1.0 × 10^13 genome copies/ml. Fluoro-Gold (Fluorochrome) and cholera toxin subunit B (CTB; List Labs) were diluted to 1% and 0.5%, respectively, with filtered water. Fentanyl and methamphetamine were obtained from the NIDA Drug Supply Program. Fentanyl (200 µg/kg), methamphetamine (1 mg/kg), and clozapine N-oxide dihydrochloride (CNO; 3 mg/kg, Hello Bio, Inc) were dissolved in 0.9% saline (10 ml/kg) and administered intraperitoneally.

### Stereotaxic Surgery

Mice were anesthetized with isoflurane (1–2%) and placed in a stereotaxic apparatus. Viral vectors or tracers were microinjected through a 34-gauge beveled needle using a syringe pump (Micro 4, World Precision Instruments) at 50 nL/min. After each injection, the needle was left in place for an additional 10 min before withdrawal. Coordinates for each targeted brain region are listed in Table 2 and are given in millimeters relative to bregma and the skull surface unless otherwise specified.

**Table 1:** Procedures and Purposes of AAVs.

| Experiment | Mouse line | Target and AAVs |
| --- | --- | --- |
| Anterograde collateral tracing | C57BL/6J | MS or LPO: AAV-retro-EF1a-Cre (200 nl);<br>SuM: AAV1-FLEX-mGFP-2A-SYP-mRuby (100 nl) |
| Retrograde tracing | C57BL/6J | LPO: 1% Fluoro-Gold (100 nl);<br>MS: 0.5% CTB (150 nl) |
| SuM-to-LPO photometry during salient stimuli | C57BL/6J | LPO: AAV-retro-EF1a-Cre (200 nl);<br>SuM: AAV9-Syn- FLEX -GCaMP7s (100-200 nl) |
| Chemogenetic manipulation | C57BL/6J | LPO: AAV-retro-EF1a-Cre (200 nl);<br>SuM: AAV9-hSyn-DIO-hM4D(Gi)-mCherry;<br>AAV9-hSyn-DIO-hM3D(Gq)-mCherry; or AAV9-hSyn-DIO-mCherry (100-200 nl) |
| Optogenetic ICSS | VGluT2-Cre | SuM: AAV9-EF1a-DIO-ChR2-EYFP, AAV9-EF1a-DIO-eNpHR3.0-EYFP, or AAV9-EF1a-DIO- EYFP (100 nl) |
| NAc DA3m during SuM-to-LPO optogenetic stimulation | VGluT2-Cre | SuM: AAV5-Syn-FLEX-rc[ChrimsonR-tdTomato] (100 nl);<br>NAc: AAV5-hSyn-GRAB-DA3m (200 nl) |

**Table 2:** Stereotaxic coordinates.

| Brain region | AP | ML | DV | Angle |
| --- | --- | --- | --- | --- |
| MS | +0.8 | 0 | -4.0 | 0° |
| LPO | 0 | +0.9 or $\pm 0.9$ | -5.1 | 0° |
| SuM | -2.7 | +0.2 or 0 | -4.6 | 0° |
| NAc | +1.1 | +0.8 | -4.5 | 0° |

For fiber photometry and optogenetics experiments, optic fibers were implanted above the target site and secured to the skull with dental cement. For fiber photometry, optic fibers (200 µm core diameter, 0.37 NA) were implanted 0.2 mm above the LPO and NAc and secured with Geristor

A and B cement (DenMat; part #4506 and #034522101). For optogenetic behavioral experiments, a 200-µm, 0.39-NA optic fiber was implanted above the right LPO. Mice were allowed 4–5 weeks for viral expression before behavioral or fiber photometry experiments. For chemogenetic experiments, a retrograde AAV-Cre vector was injected bilaterally into the LPO, and a single injection of an inhibitory (M4Di, Gi-coupled) or excitatory (M3Dq, Gq-coupled) DREADD or control vector was delivered to the midline SuM.

### Histology and Imaging

After completion of experiments, mice were deeply anesthetized and transcardially perfused with ice-cold 1× PBS followed by 10% formalin. Brains were post-fixed as appropriate, cryoprotected in 20% sucrose in PBS for 1–2 days, frozen, and coronally sectioned at 40 µm. For retrograde and viral tracing experiments, mice were perfused 1 week after tracer injection or 5 weeks after viral injection, as specified in the experiment-specific sections below.

For fluorescence immunostaining, sections were incubated in blocking buffer (PBS with 0.3% Triton X-100 and 1.5% normal donkey serum) for 30 min, followed by overnight incubation with primary antibodies diluted in blocking buffer. Sections were then washed in PBS and incubated with secondary antibodies in blocking buffer for 2 h. After a final series of PBS washes, sections were mounted on gelatin-coated slides with ProLong Diamond mounting medium (Thermo Fisher Scientific, USA). For retrograde tracing experiments, CTB and Fluoro-Gold were detected by immunostaining. Sections were incubated overnight with goat anti-CTB antibody (1:5000; List Biological Laboratories) and rabbit anti-Fluoro-Gold antibody (1:100; MilliporeSigma).

After PBS washes, sections were incubated for 4 h with donkey anti-rabbit Alexa Fluor 488 secondary antibody (1:1000; Jackson ImmunoResearch Labs, USA) and donkey anti-goat Cy3 secondary antibody (1:1000; Jackson ImmunoResearch Labs, USA). For collateral projections, fiber photometry and DREADD experiments, Cre was detected using mouse anti-Cre primary antibody (1:1000; Millipore Sigma, USA) and donkey anti-mouse-Cy5(collateral projections and fiber photometry) or A488 (DREADD) secondary antibody (1:1000; Jackson ImmunoResearch Labs, USA).

Low-magnification images were acquired with a BZ-X710 microscope (Keyence, Osaka, Japan); high-magnification images were acquired with an FV1000 (Olympus, Japan) or Zeiss LSM880 confocal microscope (Zeiss, Munich, Germany), as appropriate. Injection sites, viral expression, tracer deposition, optic-fiber placement, and fluorescent labeling were verified histologically. Animals with mistargeted injections or misplaced fibers were excluded according to predefined anatomical criteria.

### Fiber Photometry

A fiber photometry system (Doric Lenses, Quebec, Canada) with a dual-wavelength LED driver was used. Excitation light consisted of sinusoidal waveforms at 465 nm (531 Hz) for GCaMP and 405 nm (208 Hz) for the isosbestic control. Light was transmitted through a fluorescence mini-cube and patch cables (0.48 NA, 400 µm core) to the implanted optic fiber (0.37 NA, 200 µm core). Emitted fluorescence (465 nm for GCaMP and DA3m, 405 nm for the control channel) was collected through the same fiber assembly and detected by photoreceivers via bandpass filters and beamsplitters. Signals were acquired at 1,200 Hz using Doric Neuroscience Studio software and synchronized with stimulation events.

Fiber photometry data were analyzed using custom Python scripts. GCaMP, DA3m and control-channel signals were binned into 1-min epochs. The control channel was linearly fitted to the GCaMP signal, and ΔF/F was calculated as:

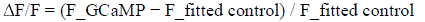

### Collateral Projections of SuM**→**MS and SuM**→**LPO Neurons

To examine collateral projections of MS– and LPO-projecting SuM neurons, we injected AAV-retro-EF1a-Cre into the MS or LPO and AAV1-FLEX-mGFP-2A-SYP-mRuby into the SuM of C57BL/6J mice. Viral vectors were infused at 50 nL/min as described above. Five weeks after surgery, mice were perfused and brains were processed for fluorescence imaging. mGFP signal was used to identify Cre-dependent somatic and axonal labeling, and SYP-mRuby signal was used to evaluate putative synaptic terminal labeling in downstream regions. Low-magnification images were acquired with the BZ-X710 (Keyence). A total of 9 mice (4 male, 5 female) were used for this experiment.

### Ratio of Collateral Projections Between MS– and LPO-Projecting SuM Neurons

To visualize SuM neurons projecting to the MS, LPO, or both targets, we injected 1% Fluoro-Gold (100 nL; FG, Fluorochrome, Colorado, USA) into the LPO and 0.5% Cholera Toxin B Subunit (150 nL; CTB, List Labs, California, USA) into the MS of 4 C57BL/6J mice. One week after tracer injection, mice were perfused and brain sections were processed for immunofluorescence staining and imaging. SuM neurons containing FG, CTB, or both tracers were identified across anatomically matched sections. Colocalization analysis was performed using Huygens Suite (Scientific Volume Imaging). Rectangular ROIs (300 × 600 µm) were placed in the right SuM across three coronal sections representing anterior, middle, and posterior SuM levels (approximately bregma −2.7 to −3.0 mm). Double-labeled neurons were counted as FG+/CTB+ cells, and counts were summarized as the number or percentage of single– and double-labeled neurons relative to the total number of retrogradely labeled SuM neurons.

### GCaMP Signal Recording in SuM Neurons Projecting to the LPO During Salient Events

To determine how SuM neurons that broadly project to the LPO and other motivation-related regions respond to threats, salient stimuli, and consumable rewards, we injected retro-AAV-Cre into the LPO and AAV-FLEX-GCaMP7s into the SuM and recorded activity in response to these stimuli.

Fiber photometry recordings were performed using the system described above. Fluorescence signals signals were acquired using Doric Neuroscience Studio software and synchronized with behavioral event timestamps generated by Med Associates programs.

For event-aligned analyses, ΔF/F signals were z-scored relative to a pre-event baseline. Mean event-aligned traces and area-under-the-curve values were extracted within predefined analysis windows for each behavioral event. Recordings were performed during the salient events described below, in the order in which they were conducted.

### Stimulus discrimination conditioning with water

Water-restricted mice were placed in a conditioning chamber equipped with a house light, an auditory cue generator, an operant (initiation) nose-poke port, and a reward nose-poke port. During the first five sessions (magazine training), mice were habituated to make a nose-poke response at the reward port (reNP), which triggered delivery of a 1-s click sound and 5 µL of water; each session lasted 60 min. Following magazine training, mice were trained on a discrimination task. A nose-poke response at the initiation port (opNP) turned off the house light and, after a 0.5-s delay, triggered presentation of one of two 2-s tones: a 20-kHz tone (CS+) was followed by 1-s activation of the reward syringe (accompanied by a brief click) beginning 1.7 s after CS+ onset, whereas a 3-kHz tone (CS−) had no consequence. The CS+/CS− tone assignment was reversed in 3 mice. Ten seconds after CS− onset, the house light was turned back on, signaling availability of the next trial, which was initiated by an opNP response. Each session lasted 60 min.

### Salient-stimulus tests

Mice were returned to ad libitum water access and placed in a pitch-dark chamber in which a 60-W lightbulb (840 lumens) was illuminated for 1 s on a variable-interval (VI) 45-s schedule, repeated 100 times. In a separate loud-sound session, a 1-s, 115-dB auditory stimulus (Tokatuker alarm siren) was presented 100 times on the same VI 45-s schedule. The order of the light and sound sessions was reversed in 3 mice to counterbalance order effects.

### Cue–footshock test

Mice underwent cue discrimination conditioning with footshock in Med Associates conditioning chambers. On CS+ trials, a 5-kHz tone was presented for 2 s, and a 0.3-s, 0.45-mA footshock was delivered 1.7 s after CS+ onset. On CS− trials, a 15-kHz tone was presented for 2 s with no footshock. CS+ and CS− trials were presented in pseudorandom order, separated by a VI 60-s schedule. The CS+/CS− tone assignment was reversed in 3 mice to counterbalance potential bias.

### Chemogenetic Manipulation of SuM Neurons Projecting to the LPO (and Their Collaterals) During Motivated Behavior

To determine whether chemogenetic activation or inhibition of SuM→LPO neurons alters behavioral responses, C57BL/6J mice received bilateral injections of a retrograde AAV-Cre vector into the LPO and a Cre-dependent M4Di(Gi)-mCherry, M3Dq(Gq)-mCherry, or mCherry control AAV into the SuM. CNO (Hello Bio, 3.0 mg/kg, i.p.) or saline (10 µL/g, i.p.) was administered 10–20 min before testing. Mice underwent the behavioral assays described below in the order presented, with at least 24 h between assays.

### Open-field test

Mice received CNO and were placed in a rectangular open-field arena (40 × 40 × 40 cm) for 20 min of free exploration. The central zone was defined as a 23.8 × 23.8 cm square centered in the arena; the remaining area was defined as the peripheral zone. Activity was recorded and analyzed using EthoVision video-tracking software.

### Water intake test

Mice underwent the water-restriction regimen and were first placed in the test chamber for two 30-min baseline sessions, during which each nose-poke response at the reward port delivered 5 µL of water. Mice then received saline and CNO (one per session, in the same chamber) and were allowed to consume water freely for 30 min in each of two subsequent sessions.

### Conditioned approach response for water

Water-restricted mice underwent Pavlovian conditioning in which a 6-s CS predicted water delivery. The CS consisted of turning off the house light and turning on cue lights at the reward port for 6 s. Five seconds after CS onset, 5 µL of water was delivered at the reward port along with a 1-s click. Mice received CNO before acquisition sessions 1–7 but not before extinction sessions 8 and 9. Each session consisted of 60 trials separated by a VI 60-s schedule.

### Conditioned place preference

After mice were returned to ad libitum water access, they underwent conditioned place preference testing in a three-compartment chamber. On day 1, mice freely explored all compartments for 15 min without CNO. Over the next 3 days, mice received saline in the morning and were confined to one compartment, and CNO or methamphetamine (1 mg/kg, IP; positive control) in the afternoon and were confined to the opposite compartment, for 30 min per session. On the test day, mice were placed without any injection treatment in the apparatus with free access to all compartments for 15 min. EthoVision XT tracked each mouse’s location to determine time spent in each compartment. Place preference was defined as the change in time spent in the CNO-paired compartment between the pre– and post-conditioning sessions.

### Fear conditioning and conditioned fear test

Fear conditioning was conducted in chambers equipped for footshock delivery and video monitoring. Mice received CNO before the conditioning session. After a 3-min baseline period, each mouse received six pairings of a white-noise CS (70 dB, 6 s) and a footshock US (0.45 mA, 1 s), with the footshock delivered 5 s after CS onset and co-terminating with the CS. CS–US pairings were separated by a VI 60-s schedule.

The following day, mice returned to the same chamber without CNO for a 7-min test session using the same CS but no footshock. During the first 3 min (no CS), freezing was assessed to measure context-driven fear; afterward, a 6-s white-noise CS was presented on a VI 60-s schedule. EthoVision XT analyzed video footage, identifying each mouse by grayscale value changes and scoring immobility as periods with pixel fluctuation below a 0.02% threshold lasting at least 0.75 s.

### Hot plate test

Nociceptive responses were assessed on a hot plate apparatus (Ugo Basile) set to 53.5 °C. After CNO administration, mice were placed on the hot plate and latency to the first nociceptive response (hind-paw withdrawal, hind-paw licking, vocalization, or jumping) was recorded, with a 30-s cutoff to prevent tissue damage (Moussawi et al., 2020). To confirm assay sensitivity, fentanyl (200 µg/kg, i.p.) was administered as a positive analgesic control. CNO and fentanyl tests were conducted on separate days.

### Tail suspension test

Tail suspension testing was performed to assess stress-coping behavior, adapted from (Ye et al., 2026). After receiving CNO, each mouse was suspended by the tail for a 6-min session in a custom tail-suspension box under continuous white noise (70–72 dB) to minimize external distractions. While the mouse rested on a platform, a 12-cm strip of adhesive tape was attached to the distal tail and secured to a horizontal bar; a lightweight plastic tube (40 × 10 × 8.5 mm, length × outer diameter × inner diameter; ∼0.9 g) was placed around the tail before suspension to reduce tail-climbing. The platform was then removed to begin the test. Sessions were video recorded, and mobility/immobility were scored using the iMOSS workflow. Primary outcomes were total immobility time and mobility/immobility bout measures over the 6-min session.

### Intracranial Self-Stimulation

VGluT2-Cre mice received AAV9-DIO-ChR2-EYFP (N = 16), AAV9-DIO-EYFP (N = 6), or AAV9-DIO-eNpHR3.0-EYFP (N = 7) into the SuM, along with an optic fiber above the right LPO.

ICSS was conducted in standard operant chambers equipped with two levers and cue lights. During the first two sessions, an active-lever press triggered a 1-s cue light with no photostimulation. In subsequent sessions, an active-lever press triggered the light cue and an 8-pulse, 25-Hz train of 473-nm photostimulation (ChR2 and EYFP groups) or 3-s continuous 575-nm photostimulation (NpHR group). Each session lasted 30 min and was repeated for up to 10 sessions.

### Photometry Procedure to Detect Dopamine Release

Mice received AAV5-DIO-ChrimsonR into the SuM and AAV5-GRAB-DA3m into the NAc, along with an optic fiber above the LPO (for stimulation) and another above the NAc (for recording). NAc dopamine signals were recorded while an 8-pulse, 25-Hz train of 638-nm red laser light (10 mW, 3-ms pulse width) was delivered to the LPO on a VI 45-s schedule. Fiber photometry recordings and data analysis were described above.

### Data Analysis and Statistics

Behavioral, histological, and photometry data were analyzed using custom Python scripts, GraphPad Prism, IBM SPSS Statistics, and other experiment-specific software.

For histological tracing, labeled neurons or fluorescence signals were quantified within predefined anatomical regions of interest. For behavioral assays, the animal was used as the statistical unit unless otherwise specified. For photometry experiments, fluorescence traces were aligned to behavioral or optical-stimulation events, normalized to a pre-event baseline, and summarized as event-aligned traces and AUC values within predefined time windows.

Data are presented as mean ± SEM unless otherwise specified. Comparisons between two groups were performed using paired or unpaired t-tests, as appropriate. Experiments involving multiple groups, sessions, treatments, or repeated observations were analyzed using one-way, two-way, or repeated-measures ANOVA, followed by appropriate post hoc comparisons when justified.

Statistical significance was set at p < 0.05.

## RESULTS

For the rest of the paper, we refer to selective projections of SuM neurons to the LPO or the MS as SuM→LPO and SuM→MS, respectively. When we refer to SuM→LPO or SuM→MS neurons and their collateral projections, we use SuM→LPO+Collateral and SuM→MS+Collateral, respectively.

### SuM**→**LPO and SuM**→**MS Neurons possess collateral projections

To characterize SuM→LPO+Collateral and SuM→MS+Collateral, we injected retrograde AAV-Cre into either the LPO or MS and delivered Cre-dependent AAV-mGFP-2A-synaptophysin-mRuby into the SuM of wild-type mice (Fig. 1A). Both SuM→LPO and SuM→MS neurons exhibited extensive collateral projections throughout the brain and showed overall similar collateralization patterns (Fig. 1B–H; Table 3). SuM→LPO neurons appeared to possess more widely distributed collateral projections than SuM→MS neurons. Notably, SuM→LPO neurons showed stronger GFP expression in the lateral habenula (LHb), whereas SuM→MS neurons exhibited more pronounced GFP labeling in hippocampal CA1 and CA2 (Fig. 1F). Although both populations projected robustly throughout the septal area adjacent to the nucleus accumbens (NAc), neither population projected directly to the NAc.

**Figure 1.**
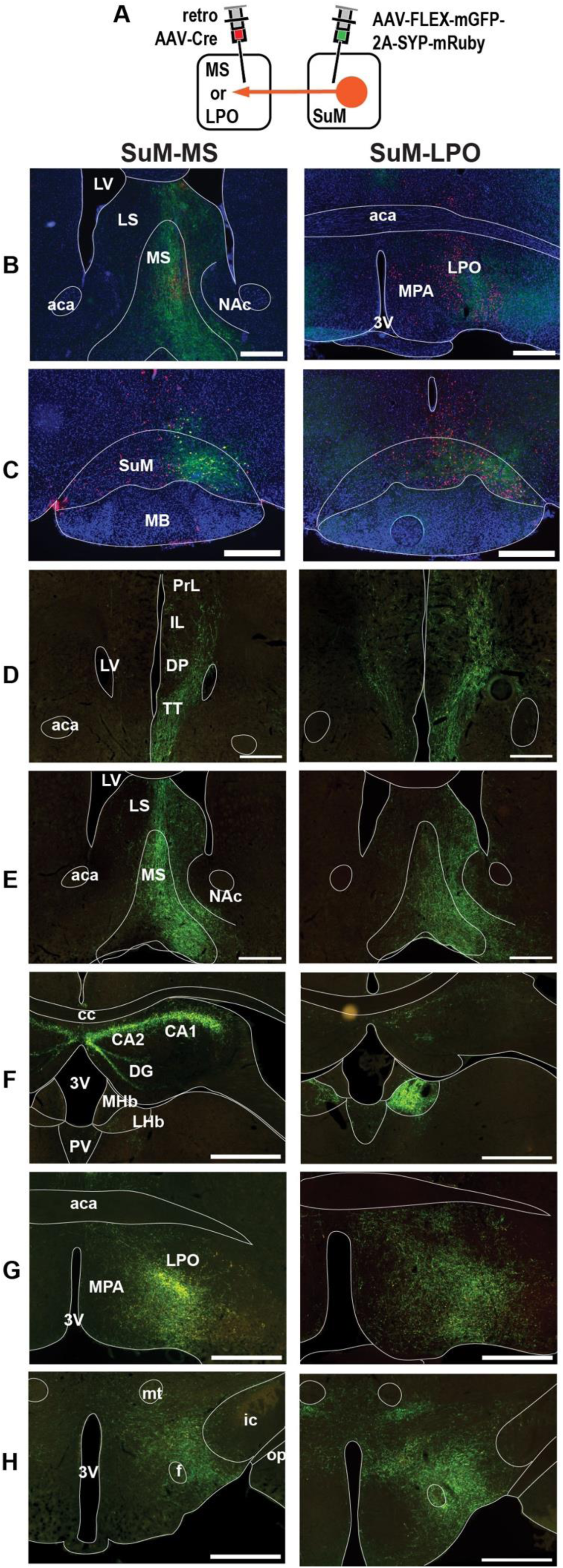
Collateral projections of MS– and LPO-projecting SuM neurons. (A) Experimental design. (B–C) Retrogradely Cre (red) labeled SuM neurons projecting to the MS or LPO, with mGFP labeling marking SuM neurons giving rise to collateralized axons. (D– H) Comparative collateral projection patterns of SuM→MS and SuM→LPO neurons across medial prefrontal cortex (D), septal region (E), hippocampal/epithalamic regions (F), preoptic region (G), and hypothalamus (H). Synaptophysin-mRuby-positive terminals were confirmed at higher magnification. Scale bars: 500 µm. Abbreviations: 3V, third ventricle; aca, anterior commissure, anterior part; CA1, hippocampal CA1 region; CA2, hippocampal CA2 region; cc, corpus callosum; DP, dorsal peduncular area; f, fornix; ic, internal capsule; IL, infralimbic cortex; LHb, lateral habenula; LPO, lateral preoptic area; LS, lateral septum; LV, lateral ventricle; MB, mammillary body; MHb, medial habenula; MPA, medial preoptic area; MS, medial septum; mt, mammillothalamic tract; och, optic chiasm; opt, optic tract; PrL, prelimbic cortex; PV, paraventricular thalamic nucleus; SuM, supramammillary nucleus; TT, tenia tecta.

**Table 3:** Collateral projections of SuM → MS and SuM → LPO.

| Group | Region | SuM-MS<br>(n=5) | SuM-LPO<br>(n=4) |
| --- | --- | --- | --- |
| Cortical region | Dorsal tenia tecta | ••• | ••• |
|  | Prelimbic and infralimbic cortex deep layer V/VI | •• | •• |
|  | Cingulate cortex 1 and 2 | • | • |
|  | Clastrum | • | • |
|  | CA1 hippocampal field | ••• | •• |
|  | CA2 hippocampal field | ••• | •• |
|  | Dentate gyrus | •• | •• |
| Basal forebrain | Vertical limb of the diagonal band | ••• | ••• |
|  | Medial septum | ••• | ••• |
|  | Lateral septum | •• | ••• |
|  | Horizontal limb of the diagonal band | ••• | ••• |
|  | Substantia innominata | ••• | ••• |
|  | Ventral pallidum | •• | •• |
| Thalamus / Epithalamus | Mediodorsal thalamic nucleus | • | • |
|  | Reuniens nucleus | • | • |
|  | Paraventricular thalamus | • | •• |
|  | Lateral thalamus | — | •• |
|  | Lateral habenula | • | ••• |
| Hypothalamus | Medial preoptic area | • | •• |
|  | Lateral preoptic area | ••• | ••• |
|  | Anterior hypothalamic area | • | • |
|  | Peduncular part of lateral hypothalamus | •• | ••• |
|  | Dorsomedial hypothalamic nucleus | •• | •• |
|  | Ventral tuberomammillary nucleus | •• | •• |
| Amygdala | Central and medial amygdala nuclei | • | •• |
|  | Amygdalohippocampal area | • | •• |
| Midbrain / Pons | Medial ventral tegmental area | • | • |
|  | Median raphe nucleus | •• | •• |
|  | Dorsal raphe nucleus | • | •• |
|  | Periaqueductal gray | • | • |
|  | Interpeduncular nucleus | — | • |
|  | Nucleus incertus | •• | •• |
|  | Laterodorsal tegmental nucleus | •• | •• |
|  | Lateral parabrachial nucleus | — | •• |

Note: The dots represent projection volume as determined by visual assessment: 3 dots indicate large, 2 dots indicate medium, 1 indicates small, and a dash denotes no detectable projection.

To quantify overlap between SuM neurons projecting to the LPO and MS, we applied a dual-tracing strategy in each mouse (N = 6): cholera toxin B (CTB) was injected into the MS and Fluoro-Gold (FG) into the LPO (Fig. 2A,B). CTB-labeled MS-projecting neurons were concentrated in the ventral SuM, extending from medial to lateral aspects (Fig. 2C). FG-labeled LPO-projecting neurons were located primarily in the medial SuM and extended dorsally and posteriorly into the ventral tegmental area (VTA).

**Figure 2.**
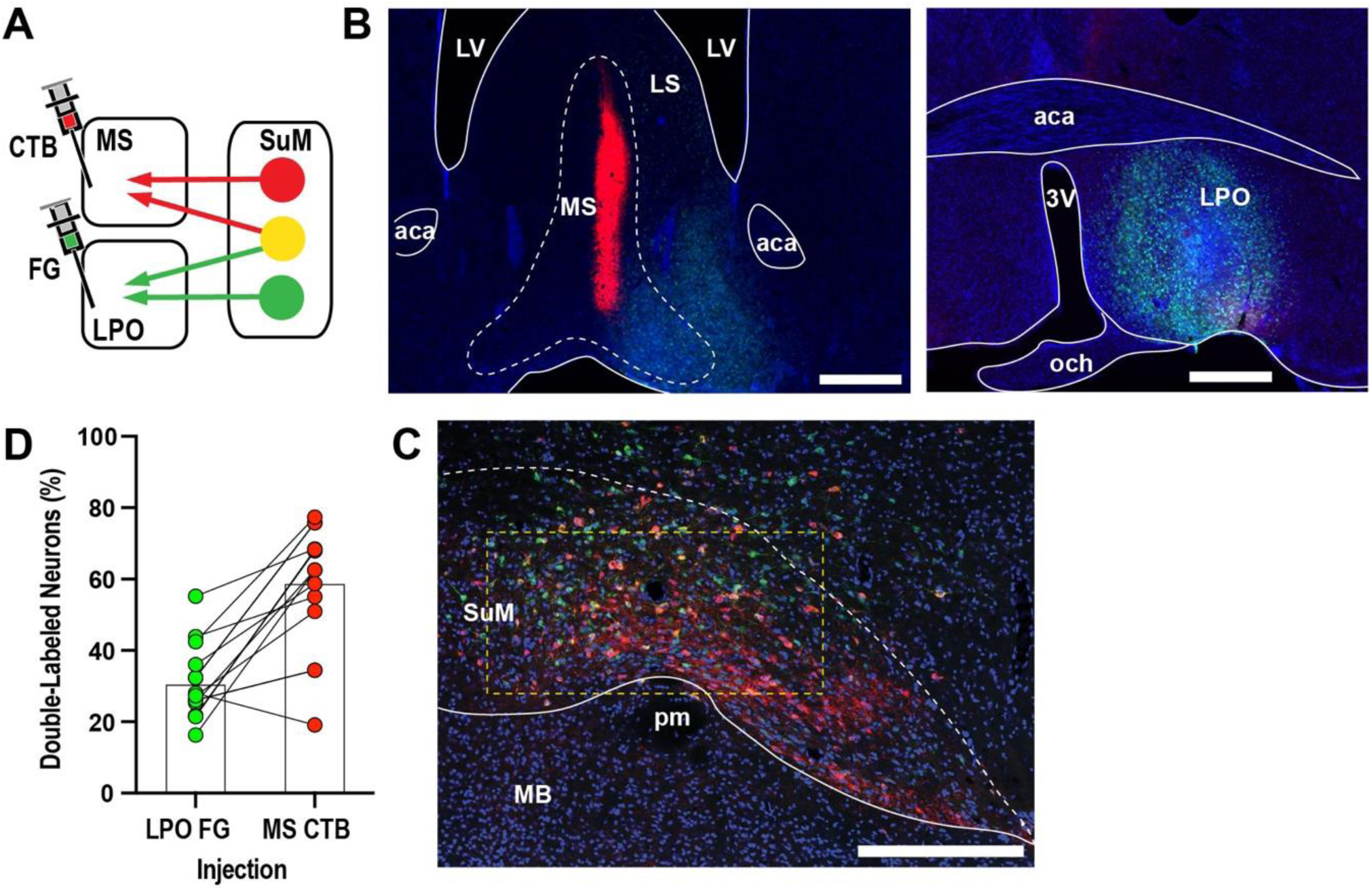
Overlap between SuM neurons projecting to the LPO and MS. (A) Schematic of dual retrograde tracer injections into MS (CTB) and LPO (FG). (B) Representative injection sites. Fig. 1 legend for abbreviations. Scale bars: 500 µm. (C) Labeled SuM neurons projecting to MS (red), LPO (green), or both (orange). The outlined ROI indicates the region quantified. (D) Percentage of double-labeled neurons.

Because the LPO tracer was delivered unilaterally, quantification was restricted to the ipsilateral SuM. Within a defined ROI (300 µm × 600 µm), dual-labeling yielded 2,100 FG-positive and 1,259 CTB-positive neurons. Of these, 30% of LPO-projecting neurons were also CTB-positive, and 59% of MS-projecting neurons were also FG-positive (Fig. 2D).

These findings indicate that most SuM→MS neurons collateralize to the LPO, whereas a smaller fraction of SuM→LPO neurons collateralize to the MS, supporting the conclusion that SuM→LPO neurons have more diverse downstream targets.

### SuM**→**LPO activity during salient positive and negative stimuli

To examine how SuM→LPO neurons distribute signals across the brain in response to salient events, we expressed Cre-dependent GCaMP7s in SuM following retro-AAV-Cre delivery to the LPO and recorded SuM→LPO activity via fiber photometry (Fig. 3A,B). Activity was measured during water ingestive behavior, bright light, loud sound, and foot shock. *SuM*→*LPO Activity Decreases During Appetitive and Consummatory Phases of Water Intake* Trials were initiated by a nose-poke at the start port (opNP), triggering a 2-second tone (Fig. 3C). The CS+ (20 kHz) predicted water delivery at the reward port (US+), whereas the CS− (3 kHz) predicted no water delivery (US−). GCaMP data were pooled across mice for sessions 1 and 9 (Fig. 3D–F), and reward and no-reward trials were analyzed separately.

**Figure 3.**
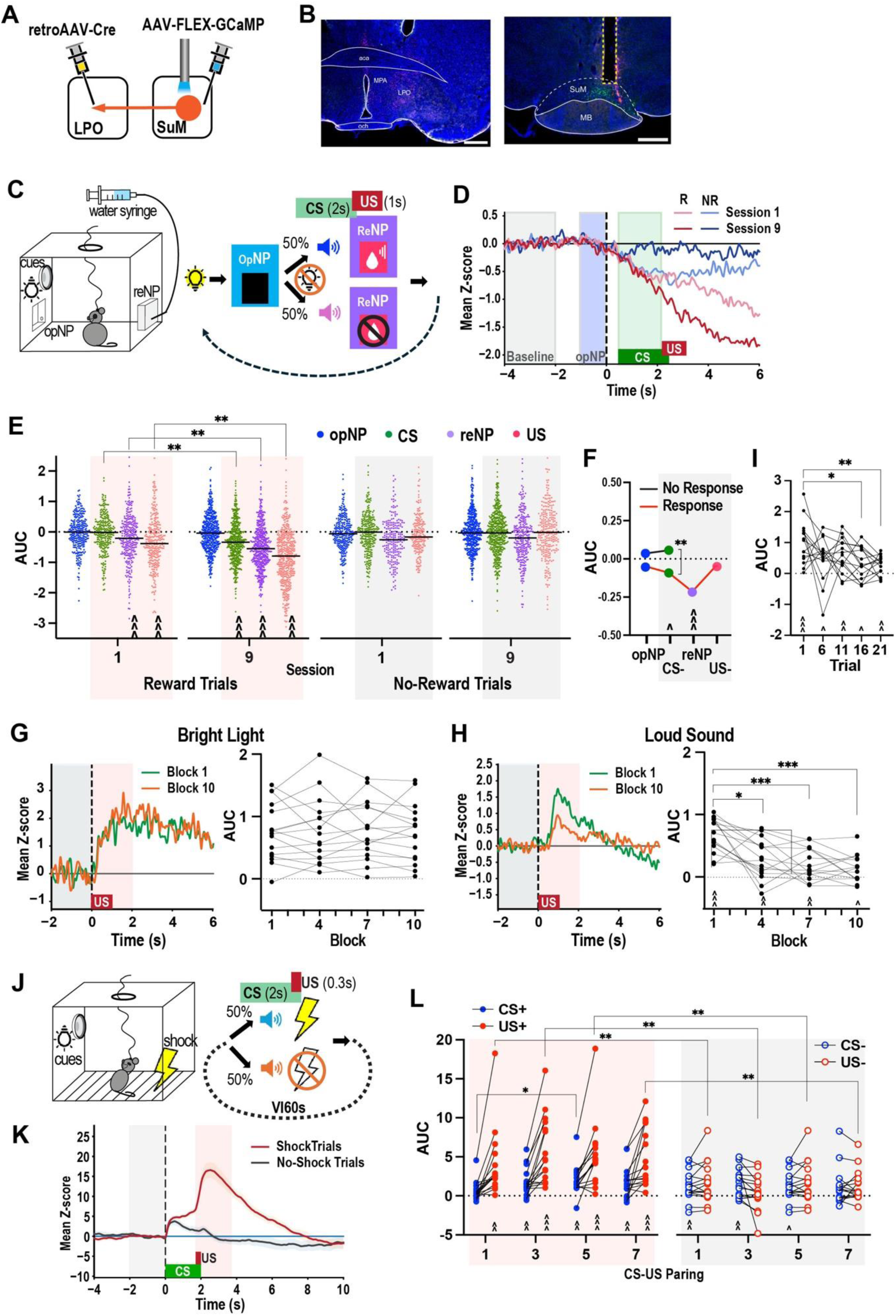
SuM → LPO GCaMP activity during motivated behavior. (A) Viral strategy for SuM→LPO photometry. (B) Example coronal sections showing LPO injections and SuM fiber placements. Fig. 1 legend for abbreviations. Scale bars: 500 µm. (C) Water-seeking task structure. (D) Representative z-scored signals for reward and no-reward trials in sessions 1 and 9. Shaded regions denote epochs used for AUC analysis. (E–F) AUC values during key behavioral epochs in reward (E) and no-reward trials (F). Sidak multiple comparisons: ^P < 0.05, ^^P < 0.01, ^^^P < 0.001 versus baseline AUC; **P < 0.01. (G–I) AUC changes following 1-s bright light or loud sound. Sidak multiple comparisons: ^P < 0.05, ^^P < 0.01, ^^^P < 0.001 versus baseline AUC; *P < 0.05, **P < 0.01, ***P < 0.001. (J) Fear-conditioning schematic. (K) Example CS-aligned traces for shock and no-shock trials. (L) AUC values across CS and US epochs in sessions 1, 3, 5, 7. Sidak multiple comparisons: ^P < 0.05, ^^P < 0.01, ^^^P < 0.001 versus baseline AUC; **P < 0.01.

### Reward trials

A mixed ANOVA with epoch (baseline, opNP, CS+, reNP, US+) and session (1, 9) produced a significant epoch-by-session interaction, F(4, 3160) = 38.57, P < 0.001. In session 1, SuM→LPO activity decreased significantly during reNP and US+, but not during opNP or CS+ (Fig. 3E). In session 9, CS+, reNP, and US+ evoked even stronger suppressions compared with both baseline and session 1. Thus, SuM→LPO neurons are suppressed during conditioned approach responses and consumption, and this suppression strengthens with training.

### No-reward trials

Because mice performed reNP on ∼60% of CS− trials, CS− trials were separated by whether reNP occurred. A three-way mixed ANOVA with epoch, session, and trial type revealed a significant trial type-by-epoch interaction, F(2, 1578) = 4.82, P = 0.008. Across sessions, CS− elicited significantly lower activity when followed by reNP compared with baseline and compared with CS− trials not followed by reNP (Fig. 3F). A second ANOVA found that reNP (but not the US− period) produced significant suppression across sessions.

Overall, SuM→LPO neurons show strong decreases in activity during reward checking and consumption, and this suppression generalizes to conditioned stimuli predicting reward. *SuM*→*LPO Neurons Increase Activity in Response to Salient Sensory Stimuli, with Stimulus-Dependent Adaptation* Mice received 100 trials of light or sound presented on a VI45-s schedule. GCaMP signals were analyzed for the 2 s before and after stimulus onset; 10-trial bins formed blocks 1, 4, 7, 10.

### Bright light

A 1-second bright light elicited strong time-locked increases in SuM→LPO activity (epoch effect: F(1, 15) = 48.45, P < 0.001) with no habituation across blocks (Fig. 3G).

### Loud sound

Loud noise also increased SuM→LPO signals, but with a significant epoch-by-block interaction, F(3, 42) = 16.38, P < 0.001 (Fig. 3H). Responses in block 1 were significantly higher than blocks 4, 7, 10.

To quantify adaptation, we examined the first 21 trials. A significant epoch-by-trial interaction emerged, F(4, 56) = 4.15, P < 0.005 (Fig. 3I): responses declined significantly by trials 16 and 21.

These findings indicate two components of SuM→LPO auditory responses: (1) a rapidly habituating component, and (2) a sustained component providing reliable responses throughout the session.

*SuM*→*LPO Neurons Increase Activity in Response to Foot Shock and Conditioned Stimuli*

Foot shock and tone cues were presented in pseudorandom order (Fig. 3J). A mixed ANOVA revealed a significant condition-by-epoch-by-trial interaction, F(6, 90) = 2.31, P = 0.040 (Fig. 3K,L). Across trials, foot shock reliably and robustly increased SuM→LPO activity.

CS+ acquired the capacity to activate SuM→LPO neurons rapidly: responses were absent on the first presentation but significant on presentations 2–4. CS− increased activity early in training, likely due to tone generalization and the sequencing of CS+ trials, but by trial 7 the CS− no longer increased SuM→LPO activity, indicating successful discrimination of CS− from CS+.

Thus, SuM→LPO neurons show strong unconditioned responses to shock and rapid learning-dependent increases to shock-predictive cues.

### Activation and inhibition of SuM**→**LPO+Collateral neurons alter motivated behaviors

We bilaterally expressed M3Dq, M4Di, or control mCherry in SuM→LPO+Collateral neurons to assess effects of pathway-wide excitation or inhibition (Fig. 4A,B). Given that SuM→LPO+Collateral neurons show opposite responses to ingestive vs threat-related stimuli, we examined behaviors across both domains.

**Figure 4.**
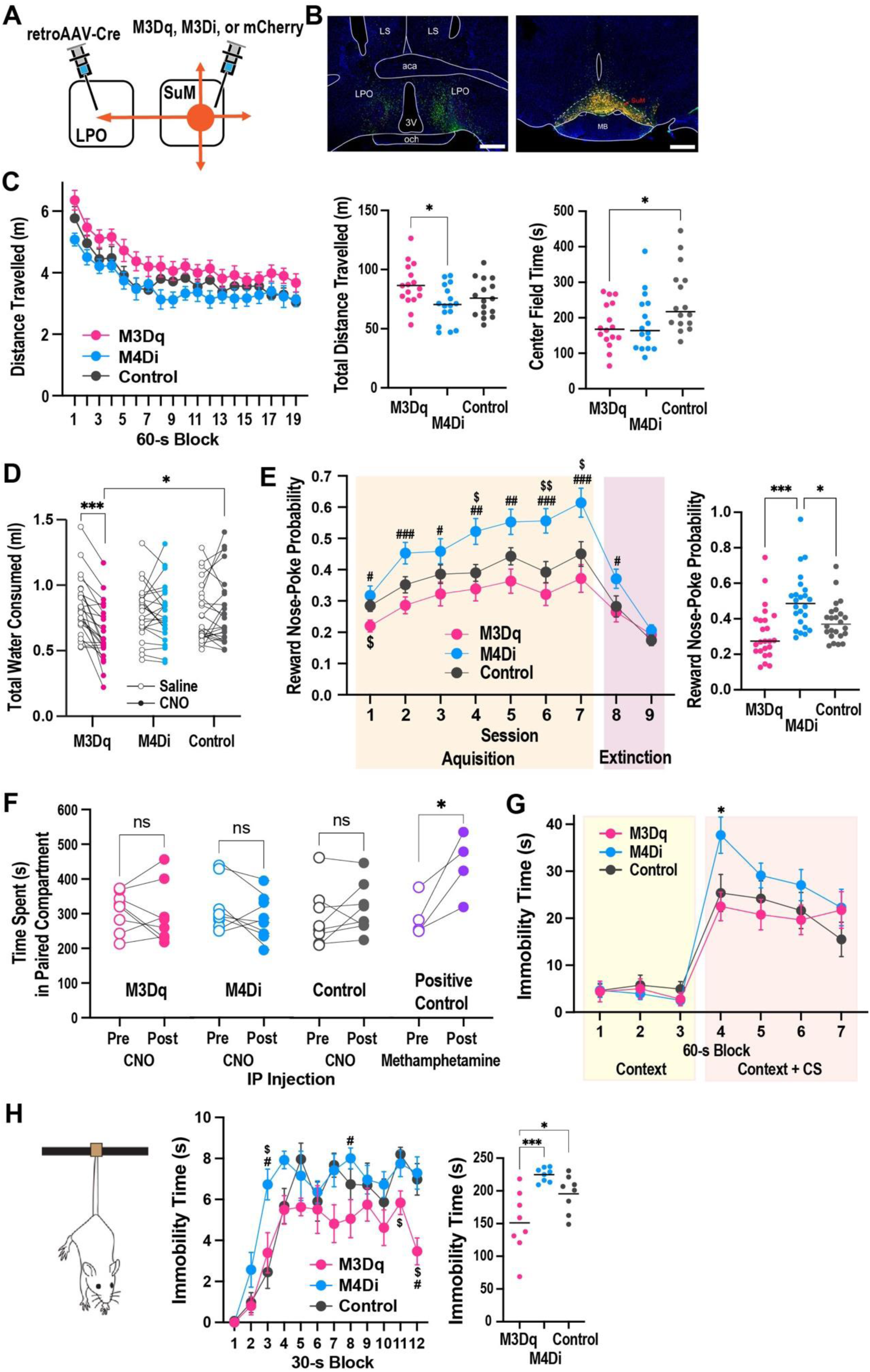
Chemogenetic manipulation of SuM → LPO+Collateral and motivated behavior. (A) Viral strategy for DREADD expression in SuM→LPO+Collateral. (B) Representative LPO and SuM labeling. See Fig. 1 legend for abbreviations. Scale bars: 500 µm. (C) Locomotion and center-time behavior in the open field. Sidak multiple comparisons: *P < 0.05. (D) Water intake after saline or CNO. Sidak multiple comparisons: *P = 0.036; ***P < 0.001. (E) CS-triggered approach probability across acquisition and extinction (left), and acquisition means across sessions 1-7 of individual mice are shown in the right. Sidak multiple comparisons: ^$^Ps < 0.05, ^$$^P < 0.01 versus control values of corresponding session; ^#^Ps < 0.05, ^##^P < 0.01, ^###^P < 0.001 versus M4Di values of corresponding session; *P < 0.05; ***P < 0.001. (F) Conditioned place preference scores. *P = 0.037, paired t test. (G) Context and CS immobility during fear-conditioning test. *P = 0.012, Tukey’s multiple comparisons. (H) Immobility during the tail-suspension test. Sidak multiple comparisons: ^$^Ps < 0.05 versus control values of corresponding session; ^#^Ps < 0.05 versus M4Di values of corresponding session; *P = 0.036; ***P < 0.001.

#### Novel Open-Field Activity

Locomotor behavior was measured in a novel open-field chamber following CNO injection. A mixed ANOVA showed a significant group effect (F(2, 45) = 3.79, P = 0.030). M3Dq mice traveled significantly farther than M4Di mice, though neither differed from controls (Fig. 4C).

Time in the center showed a significant group effect (F(2, 45) = 3.90, P = 0.028). In particular, M3Dq mice spent significantly less time in the center than controls, suggesting SuM→LPO+Collateral activation may have an anxiogenic effect.

#### Activation of SuM→LPO+Collateral Reduces Water Intake

Because SuM→LPO neurons are strongly suppressed during consumption, we hypothesized that M3Dq activation would disrupt drinking. Indeed, a significant group-by-session interaction emerged (F(2, 69) = 13.17, P < 0.001; Fig. 4D). CNO markedly reduced intake in M3Dq mice relative to saline and controls; M4Di mice showed no effect.

Thus, SuM→LPO+Collateral activation interferes with water consumption, whereas inhibition does not.

#### Inhibition Enhances Conditioned Approach Toward Water; Activation Disrupts It

Mice underwent a Pavlovian conditioning procedure in which a 6-s light predicted water delivery. The mice received CNO treatment prior to acquisition sessions 1-7, but not extinction sessions 8 and 9. A mixed ANOVA found significant group and group-by-session effects (group-by-session: F(12, 414) = 1.86, P = 0.038; group: F(2, 69) = 9.83, P < 0.001).

During acquisition, M4Di mice showed higher approach probabilities than controls in several sessions, whereas M3Dq mice showed reduced approach, especially early in training (Fig. 4E). During extinction, group differences persisted in session 8 but disappeared by session 9.

Thus, inhibition of SuM→LPO+Collateral enhances conditioned approach, whereas activation disrupts it.

#### Activation or Inhibition Does Not Produce Conditioned Place Preference or Avoidance

Affective valence effects of SuM→LPO+Collateral were examined. Because SuM→LPO neurons collateralize extensively, we anticipated that these manipulations might not produce clear CPP effects.

During the conditioning phase, the mice received saline in the morning and were confined in one compartment and received CNO or methamphetamine in the afternoon and were confined in the other compartment. A conditioned place preference test revealed that methamphetamine group, a positive control, displayed place preference (t(3) = 3.58, P = 0.037), whereas CNO manipulation of SuM→LPO+Collateral produced neither preference nor avoidance in any group (Fig. 4F).

#### Inhibition Enhances Fear Conditioning

On day 1, the mice were treated with CNO and placed in a chamber in which they received CS-footshock pairings. On day 2, the mice returned to the same chamber. The groups did not differ during the first 3 min, which was the context-only phase without CS presented. However, groups differed in immobility during the context+CS phase. A significant group-by-block effect was found, F(5.1, 113.6) = 2.46, P = 0.0365. M4Di mice showed significantly greater immobility in block 4 than M3Dq mice.

We examined whether SuM→LPO+Collateral are involved in pain perception, thereby affecting fear conditioning. While fentanyl injection significantly increased the latency to respond in hot-plate test, neither M3Dq nor M4Di had any detectable effect (Supplementary Figure 1).

These findings suggest SuM→LPO+Collateral inhibition enhances associative learning between CS and shock, whereas activation may weaken it.

#### Activation and Inhibition Bidirectionally Regulate Immobility in the Tail Suspension Test

The mice received CNO and were suspended by the tail for 6 min while immobility was measured. A mixed ANOVA revealed a significant group-by-block effect (F(22, 231) = 1.64, P = 0.039; Fig. 4H). M4Di mice entered an immobility-dominant phase (≥7–8 s of immobility per 30-s bin) ∼30 s earlier than controls, whereas M3Dq mice maintained lower immobility throughout. Thus, SuM→LPO+Collateral activity regulates coping responses during inescapable stress, with activation reducing immobility and inhibition increasing it.

### Selective SuM**→**LPO activation is reinforcing and drives NAc dopamine release

To determine whether selective SuM→LPO activation is reinforcing, Vglut2-Cre mice received Cre-dependent ChR2, NpHR, or EYFP AAV vectors in SuM and an optic fiber above the LPO (Fig. 5A,B) and underwent an optogenetics intracranial self-stimulation. During acquisition (sessions 3–7), ChR2 mice earned significantly more photostimulation than control or NpHR mice (F(2, 26) = 21.97, P < 0.001).

**Figure 5.**
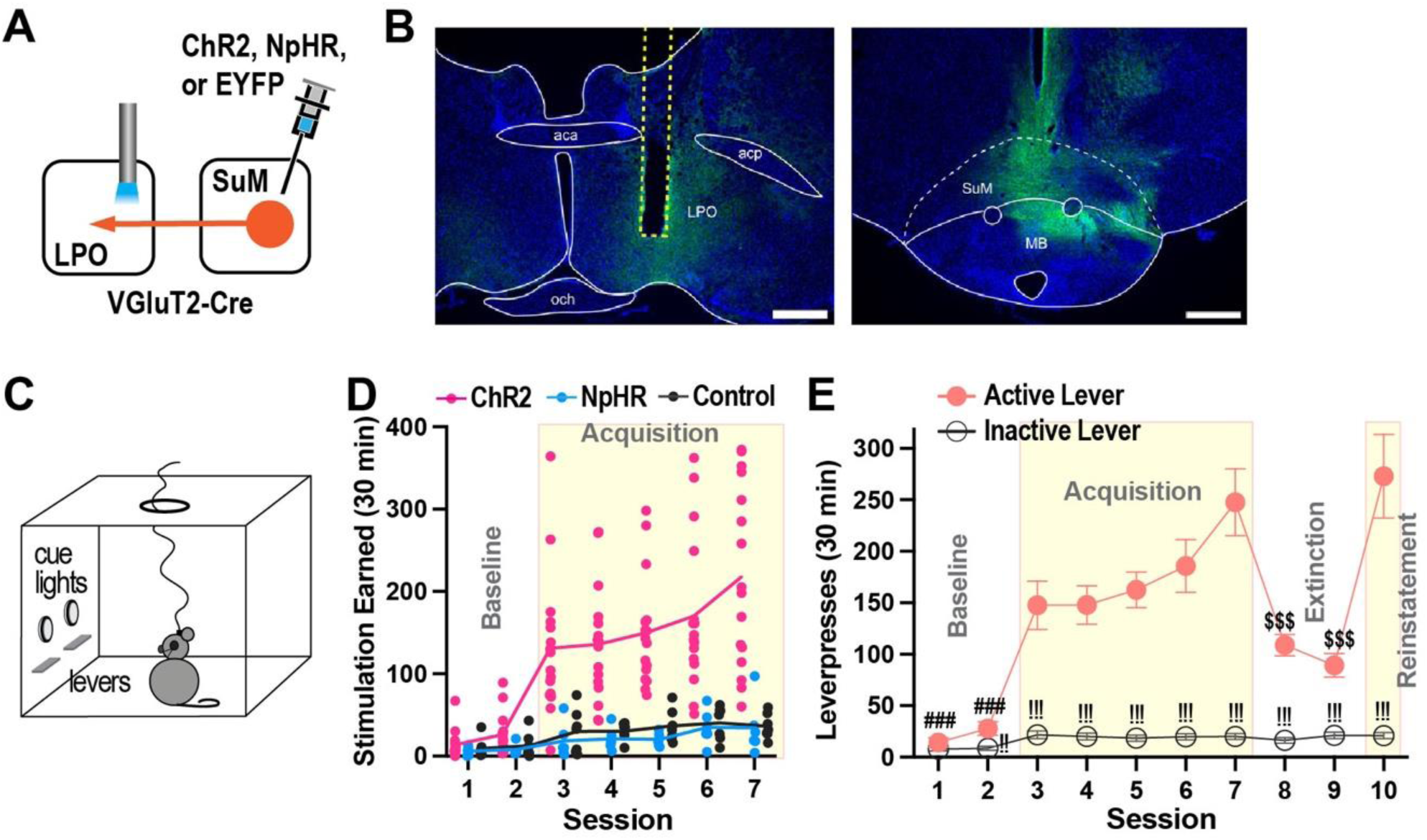
Optogenetic activation of SuM → LPO terminals reinforces approach behavior. (A) Viral injections and fiber placement in VGluT2-Cre mice. (B) Injection and fiber-track confirmation. See Fig. 1 legend for abbreviations. Scale bars: 500 µm. (C) An optogenetic intracranial self-stimulation (ICSS) chamber layout. (D) Photostimulation earned during acquisition (sessions 3–7). Lines connect session means. ChR2 mice earned significantly more stimulations than EYFP (P < 0.001) and NpHR (P < 0.001) mice, while EYFP and NpHR groups did not differ from each other (Sidak multiple comparisons). (E) Active and inactive lever presses across acquisition, extinction, and reinstatement. Sidak multiple comparisons: ^!!^Ps < 0.01, ^!!!^Ps < 0.001 versus the other lever values in corresponding sessions; ^###^Ps < 0.001 versus active-lever values of sessions 3-7; ^$$$^Ps < 0.001 versus active-lever values of sessions 7 and 10.

Lever-press analysis showed strong active-lever discrimination in sessions 2–7 and extinction-sensitive responding in sessions 8–10 (Fig. 5E). These findings demonstrate that SuM→LPO activation is positively reinforcing.

To test whether SuM→LPO stimulation drives mesolimbic dopamine release, we recorded gDA3m signals in the NAc during optogenetic stimulation of SuM→LPO terminals (Fig. 6A,B). Photostimulation produced a robust, time-locked increase in DA3m fluorescence in

**Figure 6.**
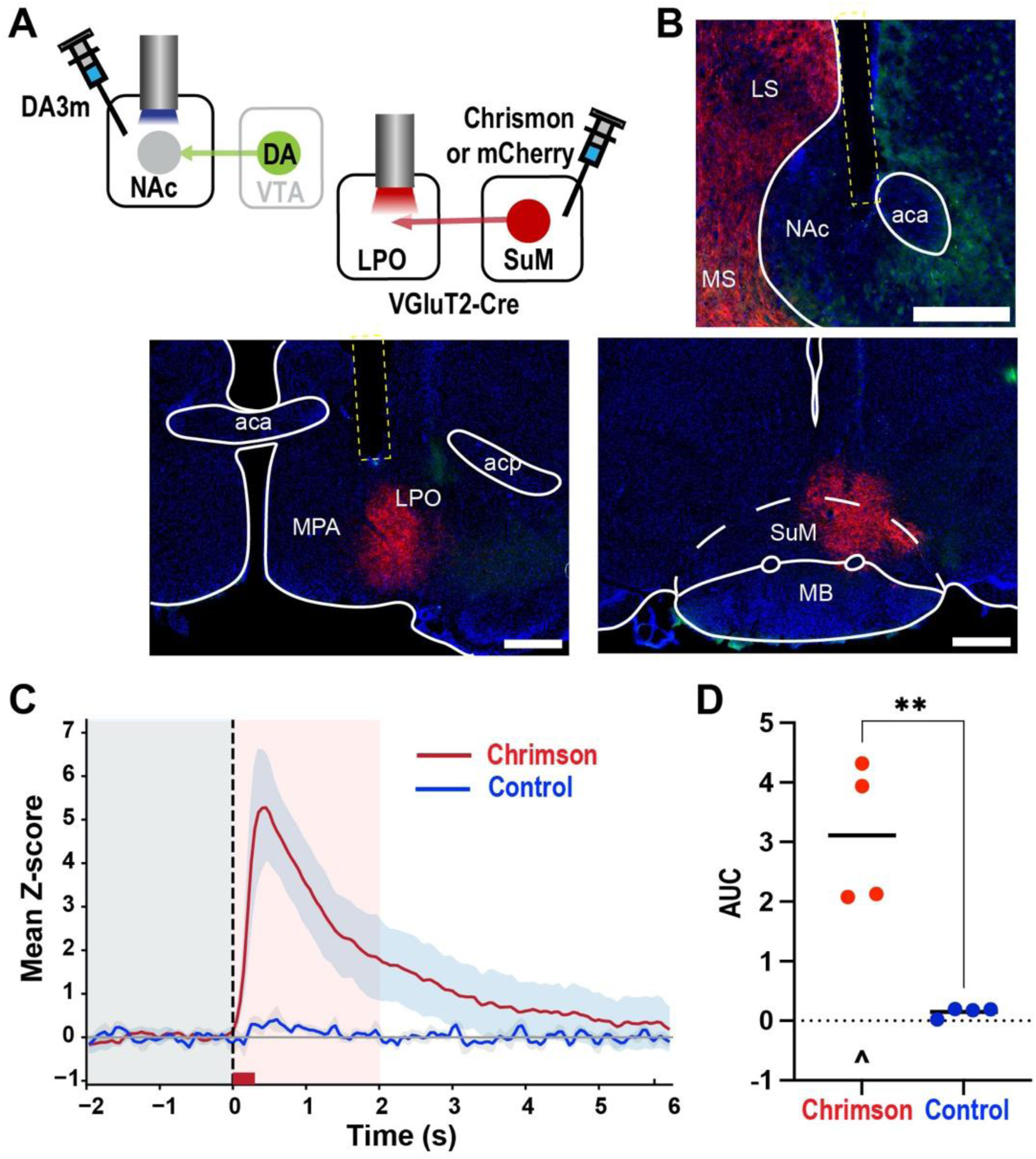
SuM → LPO terminal activation drives NAc dopamine release. (A) Viral injections for DA3m recording and Chrimson/mCherry expression. (B) Chrimson-labeled SuM→LPO terminals (red) and DA3m-expressing NAc neurons (green). See Fig. 1 legend for abbreviations. Scale bars: 500 µm. (C) Representative DA3m traces aligned to photostimulation onset (8-pulse, 25 Hz). (D) Mean AUC responses in Chrimson versus control mice. Horizontal bars show group means. Sidak multiple comparisons: **P = 0.002; ^P < 0.001 versus baseline.

ChrimsonR-expressing mice but not controls (group-by-epoch: F(1, 6) = 25.22, P = 0.002; Fig. 6C,D). Thus, SuM→LPO activation rapidly drives dopamine release in the NAc.

## DISCUSSION

The present study showed that the SuM has structural organization that could exert widespread influence over many regions through extensive collateral projections, and it identified the SuM→LPO pathway and its collaterals as circuitry through which SuM neurons can influence VTA→NAc dopamine neurons and modulate active, approach oriented actions. These findings indicate that SuM→LPO projections constitute an additional efferent route---alongside the SuM→MS pathway---that can influence VTA→NAc dopamine signaling and support active behavioral states.

### SuM**→**MS and SuM**→**LPO neurons possess collateral projections

Our anatomical experiments demonstrate that both SuM→LPO and SuM→MS neurons possess extensive collateral projections throughout the forebrain and midbrain. The overall collateral patterns were similar across populations. Some differences include that SuM→LPO collaterals were slightly more voluminous and diverse than SuM→MS collaterals and that SuM→LPO neurons have particularly strong innervation of the lateral habenula relative to SuM→MS neurons, while SuM→MS neurons showed stronger hippocampal projections. Dual labeling experiments further revealed substantial overlap between the two projection-defined populations, with most SuM→MS neurons collateralizing to the LPO, whereas fewer SuM→LPO neurons collateralized to the MS. These results indicate that information transmitted from the SuM to either LPO or MS is concurrently broadcast to multiple additional downstream nodes, enabling SuM neurons to coordinate distributed circuits involved in arousal, affect, and motor activation.

### SuM**→**LPO activity during salient positive and negative stimuli

Fiber photometry recordings revealed a clear functional organization of SuM→LPO activity across different motivational states. SuM→LPO neurons were strongly inhibited during the consummatory phase of water ingestion. This replicated our previous finding that firing rates of SuM neurons decrease during the consumption of sucrose solution (Kesner et al., 2021) and extends further by finding that SuM→LPO neurons were inhibited during conditioned approach to the reward port, and suppression strengthened across training. This indicates that SuM→LPO neurons are not simply responsive to all salient events; instead, they are specifically suppressed during appetitive and consummatory ingestion, states that require internal engagement rather than external vigilance or action. SuM→LPO neurons were also inhibited during CS− trials when mice checked the reward port, further supporting the idea that the suppression reflects the psycho-behavioral state of ingestive behavior rather than reward value.

In contrast, SuM→LPO neurons increased activity robustly in response to threats and threat-predictive cues, including unusual sensory events. Both bright light and loud noise increased SuM→LPO activity: Bright light elicited sustained increases without habituation, whereas loud noise produced an initial strong response that partially adapted over trials, suggesting a modality-selective nature of habituation and the coexistence of rapidly habituating and persistent auditory response components. SuM→LPO neurons showed strong unconditioned responses to foot shock and rapidly acquired responses to shock predictive cues, with CS+ responses emerging within a few trials. CS− responses declined once mice discriminated the two tones. Together, these findings suggest that SuM→LPO neurons respond to unconditioned stimuli and rapidly acquire the associative significance of stimuli, with their activity level modulated by external challenges.

### Activation and inhibition of SuM**→**LPO+Collateral neurons modulate motivated behaviors

Chemogenetic manipulations reveal that SuM→LPO+Collateral neurons exert opposing influences on active versus passive responses across multiple behavioral contexts. Activation of these neurons (M3Dq) increased locomotor activity and reduced immobility in the tail suspension test, consistent with promoting active coping during stress. Conversely, inhibition (M4Di) accelerated the development of immobility during inescapable tail suspension and in response to threatening cue (i.e., fear conditioning), indicating that reduced SuM output facilitates passive responding under threats.

Activation of SuM→LPO+Collateral neurons suppressed water intake and disrupted conditioned approach to water, whereas inhibition facilitated approach and strengthened conditioned appetitive response with little or no effect on consummatory response. Together with the neural activity patterns observed during photometry, these findings suggest that SuM→LPO neurons are inhibited during ingestive behaviors and that their activation can interfere with ingestive behavior.

Chemogenetic manipulations did not produce conditioned place preference or avoidance. It is unclear whether SuM→LPO+Collateral neurons are involved in generating affective valence, given that opposing effects through multiple collateral pathways could cancel out any net place preference or avoidance. Prior work shows that selective stimulation of SuM→MS pathway produces real-time place preference, whereas selective stimulation of SuM neurons projecting to the paraventricular thalamic nucleus produced real-time place avoidance (Kesner et al., 2021).

Therefore, manipulations of SuM→LPO+Collateral could have produced mixed affective effects. Given that SuM→LPO neurons send collateral projections to the lateral habenula, an established aversive hub (Shabel et al., 2012; Stamatakis and Stuber, 2012; Lecca et al., 2017), M3Dq activation may have recruited the SuM→LHb pathway. Consistent with this, our open-field data suggest that M3Dq activation is mildly anxiogenic, implying an aversive component. Moreover, the Escobedo et al study (Escobedo et al., 2024) reported that continuous photostimulation pulsed at 1 – 20 Hz over ChR2-expressing SuM→LPO+Collateral neurons induces real-time place avoidance.

### Selective SuM**→**LPO activation is reinforcing and drives NAc dopamine release

Although broad SuM→LPO+Collateral activation did not produce place preference, selective optogenetic stimulation of SuM terminals in the LPO was reinforcing in an intracranial self-stimulation task. Mice reliably worked for brief activation of SuM→LPO terminals, and this stimulation produced time-locked dopamine release in the NAc. These results identify SuM→LPO projections as a pathway that can modulate dopamine release, in addition to the previously characterized SuM→MS→VTA circuit.

Taken together with the fiber photometry results, these findings suggest that SuM neurons can mobilize active behavioral states through more than one downstream route. The SuM→MS circuit provides one established mechanism for driving VTA dopamine, but the SuM→LPO pathway represents another circuit capable of reinforcing approach behavior and modulating dopamine release.

However, it remains unanswered what precise roles parallel multiple projections play in active responses. The fiber photometry method detects population activity but not single-unit activity, and the present study manipulated SuM→LPO neurons non-selectively with respect to their extensive collateral projections. To understand the role of parallel projections, it will be necessary to examine the activity and function of SuM neurons with respect to selective, individual projection populations.

## Conclusion

This study demonstrates extensive collateral organization of SuM neurons, suggesting that SuM neurons engage multiple downstream structures in parallel to meet external challenges. The study also found that SuM→LPO+Collateral neurons can modulate active responses to external challenges, and identified the SuM→LPO pathway as a downstream pathway that can influence active, approach-oriented actions, functioning alongside the SuM→MS pathway in modulating mesolimbic dopamine. These findings significantly broaden the understanding of supramammillary function and establish the LPO as a node through which SuM neurons regulate behavioral mode, salience processing, and reinforcement.

## Ethical statement

The experimental protocol was approved by the Animal Care and Use Committee of the Intramural Research Program at NIDA and conducted in accordance with the Guide for the care and use of laboratory animals.

## Acknowledgements

The authors also acknowledge the use of the microscopy facilities provided by the NIDA Microscope Core.

