## Supplemental Information for "Supramammillary projections to the lateral preoptic area drive dopamine release and active behavior"

**SUPPLEMENTARY INFORMATION**

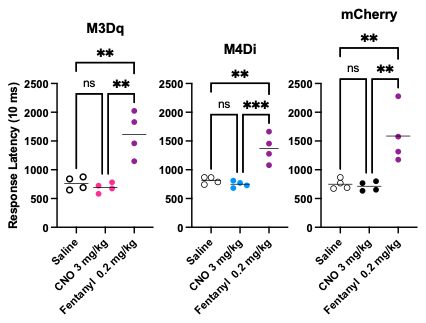

**Supplementary Figure 1.** Latency to nociceptive responses—hind-paw withdrawal, licking, vocalization, or jumping—on a 55°C hot plate was measured. One-way ANOVAs showed significant effects: M3D, (F(2, 9) = 18.51, P = 0.0006); M4D, (F(2, 9) = 20.28, P = 0.0005); mCherry control, (F(2, 9) = 11.49, P = 0.0033). Tukey’s test: **P < 0.01; ***P < 0.001.
